# Distribution, diversity and diversification from a DNA barcoding perspective: the case of *Gammarus* radiation in Europe’s oldest inland waterbody - the ancient Lake Ohrid

**DOI:** 10.1101/2023.04.18.536906

**Authors:** Tomasz Mamos, Michał Grabowski, Lidia Sworobowicz, Walter Salzburger, Sasho Trajanovski, Denis Copilaş-Ciocianu, Serena Mucciolo, Anna Wysocka

## Abstract

**Aim:** A detailed, comparative DNA-barcoding and morphospecies based overview of the vertical and horizontal distribution of Lake Ohrid endemic *Gammarus* species-flock. Re-evaluation of the origin of the species-flock dating, identification of events that putatively influenced diversification patterns in the species-flock.

**Location:** Lake Ohrid: a deep and ancient lake of tectonic origin, biosphere reserve, UNESCO World Heritage Site, located on the Macedonia/Albania border.

**Taxon:** *Gammarus* species-flock (Amphipoda, Crustacea)

**Methods:** Extensive sampling and DNA barcoding of 600 individuals were carried out. DNA sequences were analysed using species delimitation methods, haplotype network reconstructions, Bayesian molecular dating and demographic analysis. The COI-based delimitation results were validated with nuclear 28S RNA data.

**Results:** The species flock distribution has weak horizontal but clear vertical structure. The diversity across bathymetric gradients correlates with temperature and salinity; and the highest diversity with sublittoral and springs of lake’s shore. Two new MOTUs representing putatively new species are revealed and supported also by the nuclear marker. The time of flock radiation overlaps with the time of lake formation. The COI gene shows signs of positive selection and an acceleration in substitution rate. The demographic changes of the flock happened during the last ky.

**Main conclusions:** Distribution of the *Gammarus* species-flock is vertically structured, reflecting habitat zonation. Parapatric speciation as one of the mechanisms in flock’s diversification is suggested. Detection of new MOTU suggests that the flock’s diversity is still not fully revealed. Nevertheless, failure to recover three other MOTUs suggests the loss of gammarid diversity in the lake. This represents,together with the current threats to the lake ecosystem (i.e. climate changes, development of tourism), a clear call for conservation efforts. The speciation events and demographic changes within the flock relate presumably to glacial and postglacial water level changes and to colonisation of new depth ranges and the associated springs.

## 1. Introduction

From a geological perspective, most European lakes are short-lived or even ephemeral water bodies that originated at the Pleistocene/Holocene transition as a consequence of the retreat of glaciers. Ancient lakes, on the other hand, are the extant lakes that have continuously existed for more than 100 ky BP (Hampton et al., 2018; Martens, 1997;), in some cases for millions of years, meaning that they originated during or before the Pleistocene. Many of the world’s ancient lakes – for example, Lake Baikal, Lake Titicaca, Lake Matano or the Great Rift Lakes in Africa – are famous for the rate of *in situ* speciation and for spectacular examples of adaptive radiation. Despite covering less than 1% of continental surface area (Dudgeon et al., 2006), ancient lakes are hotspots of biodiversity and endemism, and natural laboratories to study evolutionary processes (Cristescu et al., 2010; Salzburger et al., 2014).

For over a century, ancient lakes have been in the focus of biodiversity-centred research. A wealth of data regarding species richness at a site (alpha-diversity), species richness variation between sites (beta-diversity), or total species diversity at the lake scale (gamma-diversity) have been accumulated (Albrecht et al., 2012; Albrecht & Wilke, 2008; Salzburger et al., 2014), while studies focusing on factors and processes determining the spatial partitioning of diversity are much less common. There is also a need for the integration of molecular and morphological data in the study of many taxa in such systems (Snoeks et al., 2000, but see Ronco et al., 2021).

While some of the world’s ancient lakes and their faunas are studied in some detail (e.g. MacDonald et al., 2005; Meier et al., 2017; Ronco et al., 2021), surprisingly little is known about the ancient lakes of Europe. It is not even fully established, which European lacustrine systems would actually qualify as ancient lakes. The only unquestionably European ancient lake, with a well-documented geological history covering the last 1.3 My, is Lake Ohrid (Wagner et al., 2014; Wagner et al., 2017; Wilke et al., 2020). This oligotrophic and karstic lake is situated in a steep-sided graben with a rift formation origin, in the South Adriatic–Ionian biogeographic region (Banarescu, 1991) on the Balkan Peninsula, at the border between North Macedonia and Albania. It has a maximum depth of 293 m, a surface area of 358 km^2^, and a volume of 55 km^3^ (Matzinger et al., 2006), which makes it one of the smallest ancient lakes in the world. Despite the small size of the lake, different types of bedrock, active tectonics (Hoffmann et al., 2010), steep-sided surrounding mountain ranges, sublacustrine and lake-shore spring fields, as well water-discharge by lake-side (Matzinger et al., 2006) determine the high habitat complexity in its basin.

The observed vertical (bathymetrical) zonation within the lake is based on the range of water temperatures, the degree of sunlight penetration, substratum, vegetation, and water movement (Radoman, 1985). The lake includes three vertical general habitat zones with two subzones, from top to bottom: (1a) an upper littoral zone with a rocky/sandy upper part (depth: 0-3 m); (1b) a lower littoral zone overgrown with *Chara* spp. (aka *Chara*-belt, depth: 3-20 m); (2a) an upper sublittoral zone (aka “shell zone” with abundant population of *Dreissena* sp. (both alive colonies and empty shells, depth: 20-35 m); (2b) a lower sandy/silty sublittoral zone without shells (depth: 35-50 m); and (3) a profundal muddy zone (depth 50-bottom). However, these zones are not evenly distributed around Lake Ohrid due to vertical diversification with a steep eastern and southeastern rocky part and, contrasting to it, a more shallow, sandy, northern part of the lake (Albrecht & Wilke, 2008).

The spatial distribution of the endemic biodiversity in Lake Ohrid is still not fully understood, especially in the context of evolutionary processes that have shaped it (Hauffe et al., 2011). Previous studies showed that, taking into account its surface, the lake is characterised by a high rate of endemism (36%) compared to other ancient lakes (Albrecht & Wilke, 2008), such as for example the much bigger Lake Victoria (31.3 %). So far, nearly 350 endemic species have been described from Lake Ohrid, of which at least 188 are animals (Albrecht & Wilke, 2008; Hauffe et al., 2015; Pešič, 2015; Stelbrink et al., 2016; Stocchino et al., 2013).

Lake Ohrid region was designated a UNESCO World Heritage Site for its natural values in 1979 and for its cultural heritage values a year later (Centre, 2022). However, the region suffers from direct and indirect human related factors (Kostoski et al., 2010; Trajanovski et al., 2019), like for example effects of climate change (Matzinger et al., 2006, 2007). As a result, the Reactive Monitoring Mission of UNESCO (Decision of the Committee for the world’s heritage of UNESCO 41-COM 7B.34), in its report on the state of the natural and cultural heritage of the Ohrid region, emphasises the problem with accumulating anthropogenic pressure due to the illegal urbanisation, no adequate waste management, and uncontrolled usurpation of the shoreline natural habitats as serious problems that could potentially affect the uniqueness and peculiarity of the Lake Ohrid ecosystem. All this caused that in 2019 and 2021 efforts were taken to change the status of Lake Ohrid into World Heritage Sites in Danger. Therefore, it is of utmost importance to properly identify and catalogue the endemic biodiversity of Lake Ohrid.

Among the endemic species in Lake Ohrid, gammarids (Crustacea, Amphipoda) are one of the most outstanding examples, with 12 of the 13 taxa being endemic (Grabowski et al., 2017; Wysocka et al., 2013). This *Gammarus* species-flock originated from the *Gammarus balcanicus* species group (Karaman & Pinkster, 1987), native in mountain springs and streams of Southern, Central and Eastern Europe. The *G. balcanicus* group is known for high molecular diversity as well as for lack of clear taxonomic features, resulting in a huge cryptic diversity (Mamos et al., 2014; Mamos et al., 2016; Wysocka et al., 2014). Addressing such ‘hidden diversity’ was possible primarily thanks to DNA barcoding based on the standard 5’ fragment of the mitochondrial cytochrome oxidase subunit I gene (COI), designed to provide rapid and automatable species identification (Hebert et al., 2003). On the other side, DNA barcodes aid in identification of animals with excessive phenotypic polymorphism, where different morphotypes appear to be conspecific or even belong to the same COI lineage (Osikowski et al., 2018). Detection and delimitation of new species based on single-locus data, such as COI, with rapidly developing bioinformatic tools, have become a common and fast approach in biodiversity assessments, particularly in case of poorly known ecosystems or animal groups lacking proper taxonomic expertise (e.g. Gadawski et al. 2022). DNA barcoding is a methodologically sound and conceptually simple method offering insights into diversification processes across many branches of the Tree of Life. Especially in the face of taxonomic impediment, when using the fast and handy method for preliminary estimation of phylogenetic lineage diversity, to highlight taxa that should be prioritised for a detailed study, is sorely needed. The COI marker proved to be divergent enough to reveal inter- and intra-specific diversity in gammarids (Csapó et al., 2020; Wattier et al., 2020) and often show congruence for interspecies diversity with 28S rDNA marker (Mamos et al., 2021; Weber et al., 2022). DNA barcoding also proved to be irreplaceable with respect to the *Gammarus* species-flock from Lake Ohrid proper, where two species were described solely based on their molecular distinctiveness (Grabowski et al., 2017).

The history of Lake Ohrid is well-studied, thanks to the SCOPSCO drilling project (Wagner et al., 2014, 2017). In particular, the diatom fossil record of Lake Ohrid provided important information on speciation and extinction patterns throughout its geological history (Jovanovska et al., 2022; Wilke et al., 2020). However, the actual age of other species-flocks in Lake Ohrid, as well as their (*in situ*) diversification patterns, has not yet been properly studied. So far, the *Gammarus* radiation was dated on the basis of a single secondary calibration point from distantly related lineages of gammarids and arbitrary constraints on terminal branches (Wysocka et al., 2013). A first meta-analysis of radiation time for invertebrates endemic to Lake Ohrid, including *Gammarus* flock, was done by Stelbrink et al., (2020). This work provided important insight into evolution of invertebrate species-flocks in the light of the lake’s limnological history. However, calibration was done on general protostomian substitution rate for COI, which may not be proper for some of the taxa. Initial hypotheses about the diversification dynamics of *Gammarus* flock through time in Lake Ohrid were proposed, including e.g. the impact of water fluctuation in the lake on the demography of members of flock, but these assumptions have never been formally tested. Furthermore, neither the horizontal nor the vertical distribution of particular species belonging to the flock has been examined so far. However, such data in Europe’s oldest lake might provide important insights into the evolutionary and ecological processes behind the phenomenon of adaptive radiation in insular aquatic environments. In addition, the proper recognition of biodiversity and its distribution in this UNESCO World Heritage Site would be crucial for the planning of future conservation actions.

The goal of our work, that is based on the extensive sampling across the entire depth profile of Lake Ohrid along several spatially distant transects, is to provide detailed information on the vertical and horizontal distribution of the local endemic *Gammarus* species-flock. We achieved this by combining the morphology-based taxonomical data on the species diversity with the DNA-barcoding data, offering fast and cross-sectional insight in the potential cryptic diversification signal within conventionally recognised taxa. In addition, we examined the usability of COI, validated by 28S rDNA marker, in dating the origin of the species-flock, as well as to identify events that putatively influenced diversification patterns in the species-flock. Generally, from the biogeographical perspective, zonation of spatially confined habitats (habitat discontinuity) can be seen as a driving force behind intralacustrine speciation, particularly in case of the relatively low-mobile benthic organisms (Brooks, 1950). On the other hand, long-term presence of habitat patches inhabited by closely related species competing for resources may promote hybridisation (Mallet, 2007; Seehausen, 2004) and slow down the achievement of a state of reciprocal monophyly in the radiating species. Taking it into account, we verify if: : (*i*) the distribution of the different morphologically determined species and COI lineages within the flock reflects the lake habitat zonation; (*ii*) the sublittoral zone being likely is the most stable over time and connecting different habitats features the highest species-richness and genetic diversity; (*iii*) the radiation of the flock coincides in time with the establishment of permanent deep-water conditions and habitat zonation in Lake Ohrid; and (*iv*) the main factor influencing demographic changes in the *Gammarus* species-flock in Lake Ohrid are water-level fluctuations caused by Pleistocene glaciations and postglacial water level increment.

## 2. Methods

### 2.1 Sampling

The new material presented in this study was gathered during scientific expeditions to Lake Ohrid in 2019 and 2022, performed in collaboration with the Hydro-Biological Institute, Lake Ohrid. Samples were collected (*i*) from a trawler boat with a fine-mesh dredge across the transect line, perpendicular to the shore, transects; (*ii*) with the Zawal’s light traps (Zawal, 2018) and artificial spruce as trap, from various depths of the lake; (*iii*) with benthic hand-nets from the shallow littoral (depth 0.25-0.5 m); and/or (*iv*) with benthic hand-nets from the springs located along the lake shoreline (Fig. 1, BOLD DS-OHLGADD). The gammarids were fixed in 96% ethanol directly after collection or after morphological examination done on land. The taxonomic assignment to a species belonging to the *Gammarus* species-flock of Lake Ohrid was determined using the morphological key provided in Grabowski et al., (2017). We also included previously published specimen material from Wysocka et al. (2013), Wysocka et al. (2014), and Grabowski et al. (2017). In case of part of the material collected in 2022, the whole bodies were ground in order to quickly obtain high quality nucleic acids and morphological examination was not performed. A list of specimens with metadata is available in DS-OHLGADD (doi: provided after acceptance). As an additional proxy, we recorded temperature and relative salinity (through speed of sound in water) in the lake water column from depths 0 m down to 250 m in May 2022, using a Valeport SWIFTplus probe.

**Fig. 1.**
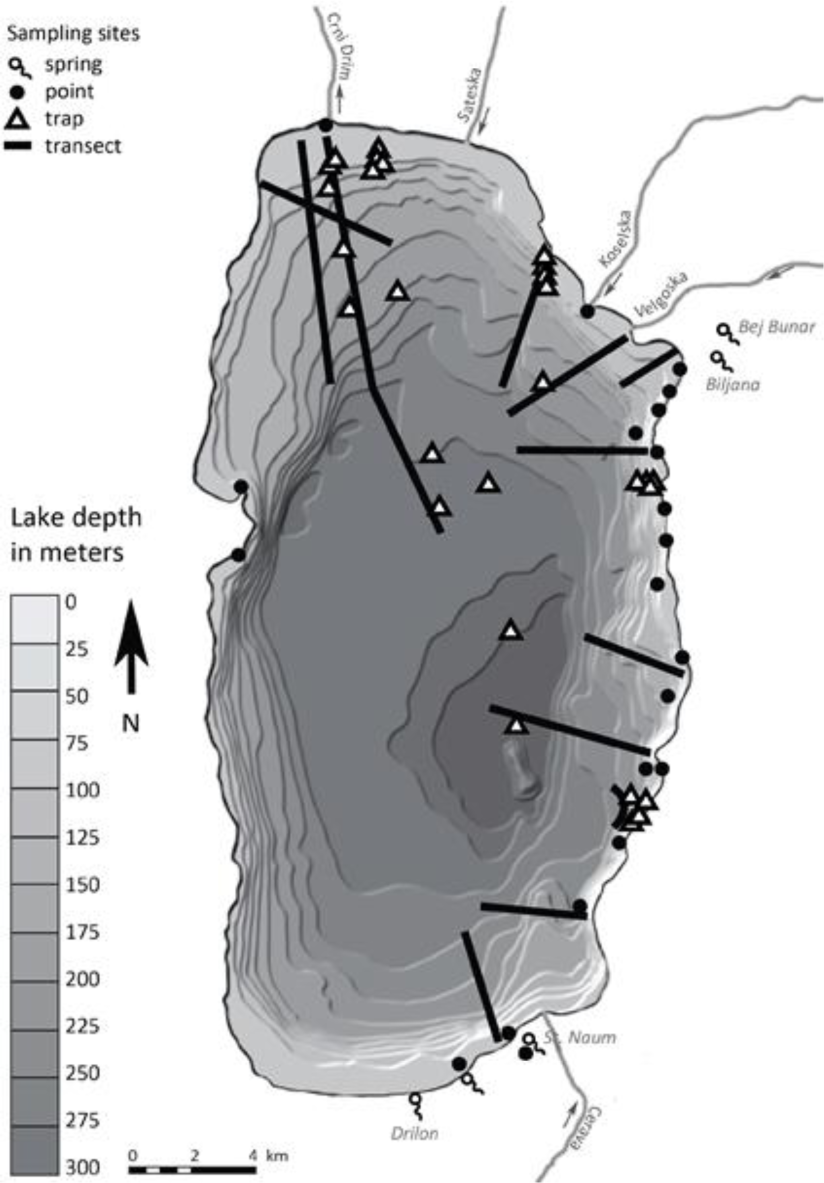
Sampling localities in Lake Ohrid and adjacent springs used in this study. Map modified from Reed et al. (2010).

### 2.2 Molecular data acquisition and initial analysis

After initial morphological identification, whole specimens were homogenised (FastPrep-24; MP Biomedicals), and total nucleic acids were isolated using the Direct-zol RNA kit (Zymo) according to the manufacturer’s protocol. The DNA subsample was aliquoted prior DNA digestion and cleaned through ethanol precipitation, then suspended in the TE buffer. The RNA was stored for future research. The COI marker (mtDNA) was PCR-amplified using the primer pair LCO1490-JJ and HCO2198-JJ (Astrin & Stüben, 2008). Additionally, the 28S rDNA nuclear marker was amplified on selected individuals, to validate the Molecular Operational Units (MOTU’s) defined based on COI and for reconstruction of dated phylogeny. Primer pairs used for the 28S marker were: 28F, 28R, 28S-700F and 28S-1000R (Hou et al., 2007).

Amplification, visualisation, and enzymatic cleaning of the PCR products followed the protocols described in Mamos et al. (2016). Sanger sequencing of PCR products was performed at Macrogen Europe using BigDye™ Terminator v3.1 on the 3730xl DNA Analyzer.

The obtained raw sequence data were examined for contamination through blast searches (Altschul, 1990) against the nucleotide database. Sequences were assembled in Geneious R11 (https://www.geneious.com) and aligned using MAFFT (Katoh & Standley, 2013), using default settings. All COI sequences were translated into amino acids in order to check for premature stop codons to exclude putative pseudogenes, however, none were detected. All newly generated sequences (COI and 28S) were deposited in the GenBank and BOLD dataset DS-OHLGADD . The data set was supplemented with already published sequences available in GenBank . Altogether, our data set consisted of 600 specimens, all with their respective COI marker sequences and 40 sequences of 28S rDNA marker. Prior to all analysis, the COI alignment was trimmed to final 575 bp in order to remove primers low quality ends, and to obtain equal length of all analysed sequences without unknown terminal parts, as it is required by some of the performed analysis. The COI haplotypes were determined using DnaSP 6 (Rozas et al., 2017).

In order to identify putative molecular operational taxonomic units (MOTUs), two distance based and two phylogeny based species delimitation methods were used. For the distance methods, all COI sequences were automatically assigned a Barcode Index Number (BIN) in BOLD. The BIN system clusters barcode sequences algorithmically through refined single linkage analysis in four steps, to calculate MOTUs showing high concordance to species. The four steps included: sequence alignment, generation of initial MOTU boundaries based on single linkage clustering, evaluation of opportunities for refinement of the boundaries using Markov clustering and finally, selection of the optimal partitions for MOTUs based on the Silhouette index (Ratnasingham & Hebert, 2013). Additionally, we run ASAP – assemble species by automatic partitioning (Puillandre et al., 2021) – on a dataset of all COI sequences using the K2p distance model (Kimura, 1980). For the phylogeny based delimitation method, we reconstructed a Maximum Likelihood tree in RAxML 8.2.12 (Stamatakis, 2014). The substitution model (TPM3uf+I+G4) was selected using ModelTest-NG (Darriba et al., 2020) and the bootstrapping function set to autoMRE. The tree was used as proxy for Bayesian implementation of the Poison Tree Processor (bPTP, Zhang et al., 2013), performed on the bPTP web server (available at https://species.h-its.org) and the multi-rate PTP (mPTP, Kapli et al., 2017) which implements MCMC sampling that provides a fast and comprehensive evaluation of the inferred delimitation. Both methods were run with 500,000 iterations of MCMC and 10 % burn-in. The p-distance and the K2p-distance were calculated in Mega 11 (Tamura et al., 2021). In order to visualise molecular distances and relationships between haplotypes, phylogenetic network was constructed in SplitsTree 4 (Huson & Bryant, 2006) for COI haplotypes using the Neighbour-Net (Bryant & Moulton, 2004) algorithm with uncorrected p-distance and haplotype network was reconstructed for 28S using the TCS algorithm (Clement et al., 2000) in PopArt v. 1.7 (Leigh et al., 2015). The 28S sequences were trimmed in this analysis to 777bp and sequences containing unknown nucleotides were removed.

### 2.3 Community and ecology analysis

The available physical data (depth and relative salinity) and number of MOTUs were averaged for every 10 m intervals. Data were visualised as boxplots (with median, quartile range and outliers) and basic statistics such as mean and standard deviation were calculated in R (R Core Team, 2022). The same bathymetric partitioning was used to calculate means and standard deviations for physical factors and the number of MOTUs. To explore possible associations between the number and nature of MOTUs and depth, temperature and relative salinity, the correlation coefficients and their p-values were calculated. Because the physical data did not fit a normal distribution, we performed Spearman correlation analyses. Additionally, to test if correlations are significant despite violating the assumption of neutral diversity, we applied Pearson correlations. Correlation calculations were performed with corr.test::psych function, together with visualisations, in R. Statistics of DNA sequence polymorphism such as haplotype diversity (H_d_) and nucleotide diversity (π) were calculated using DnaSP 6 (Rozas et al., 2017). These statistics were calculated for sequences from springs and three major habitat zones – littoral, sublittoral, profundal – following the zonation proposed by Albrecht & Wilke (2008) and Radoman (1985).

In order to assess if the community composition is structured by habitat zones, we run an analysis of variance using distance matrices (permutational MANOVA) with adonis2::vegan function in R (Anderson, 2001). The every 10 m grouping was used for generating the community dataset. Because the profundal zone spread over a wide depth and lower number of samples were collected not every 10 m, we collapsed it into 4 groups with similar abundances as in groups from littoral and sublittoral. We calculated two data distance matrices with vegdist::vegan, one using log transformed MOTUs abundance (Euclidean method), the other using only the presence/absence matrix (Jaccard method). Additionally, we run the variance analysis on two datasets with and without the spring communities. The permutation test was performed with 999 replicates. To visualise the relationships between communities, including springs, we performed non-metric multidimensional scaling analysis (NMDS) with metaMDS::vegan function. The stress parameter was recorded, and Shepard plot was used to assess data loss during dimension scaling and the solution converged. Using the function envfit::vegan we have fitted the species abundance to the ordination, checking which species significantly influenced the ordination.

### 2.4 Analysis of population structure

To explore the genetic connectivity and population structure of gammarids across the depth gradient in Lake Ohrid, analysis of molecular variance (AMOVA) and F_st_ were calculated for the three MOTUs featuring the widest vertical range. Here, every 10 m depth interval was treated as one site and the sites were then grouped according to the conventionally accepted three habitat zones (see above).

To explore the patterns of gene exchange between species from the springs and from the habitat zones in the lake, we applied a Bayesian coalescence model implemented in MIGRATE-N 4.4 (Beerli et al., 2019) that infers the migration rate (M = m/μ, where m is the immigration rate per generation). The analyses were run for the three MOTUs with the greatest distribution range (LYCH-PAR-OCH, STA-LYCH, SOL) and for which we had data on more than 10 specimens per habitat zone. Initial runs were performed in order to assess priors and MCMC settings. To perform coalescent migration analyses, the HKY (Hasegawa et al., 1985) model was applied. In most analyses, default priors were used except for MOTU LYCH-PAR-OCH, where the upper *prior* boundary of migration parameter was set to 50k, after examining the posterior distribution of the initial runs. The MCMC simulation was run with one long chain with 50000 recorded genealogies and a 10% burn-in, a sampling increment of 500, and a heating scheme with four chains and temperatures (1.0, 1.5, 3.0, 1M). Effective sampling sizes (ESS) of the parameters returned with high values (>1000). To test the hypothesis that the different habitat zones act as barriers for gene flow, we designed different models representing various hypothesis: (*i*) a matrix model with separate populations allowing migration between all populations; (*ii*) a model with one panmictic population across all habitat zones; and (*iii*) a model assuming an asymmetric migration from lake to springs in the case of MOTU LYCH-PAR-OCH. After runs of different models, the log-marginal-likelihood of the raw thermodynamic score and Bezier approximate scores were examined.

### 2.5 Phylogeny reconstruction and molecular clock dating

Prior to molecular dating, in order to investigate putative conflicts in phylogenies, we run phylogeny reconstruction for COI and 28S markers separately, using specimens for which we have generated 28S rDNA. For phylogeny reconstruction we used BEAST 2.6.2 (Bouckaert et al., 2019), with substitution model set through bModelTest (Bouckaert & Drummond, 2017, resulting models with highest posterior probability for this and following Bayesian analysis given in Table S2.1), Birth-Death tree prior and without setting molecular clock rate (default = 1). Two MCMC runs, each with 20,000,000 generations and a sample frequency every 2,000 generations were applied. All runs and parameters were examined and proved to have ESS>200. The runs were combined, and final trees annotated using Logcombiner and TreeAnnotator, respectively (Bouckaert et al., 2019). The resulting trees were plotted using cophyloplot::ape function in R in order to show links between sequences coming from different markers but the same specimens.

In order to interpret the time-line of diversification of the *Gammarus* species-flock in Lake Ohrid, a molecular clock method was employed. To this end, a Bayesian time-calibrated phylogeny reconstruction was performed with BEAST 2.6.2. For the reconstruction we used one specimen per MOTU and the COI and 28S sequences of the endemic species as well as *G. sketi,* its sister lineage. The calibration was based on the fossil remains of amphipods preserved in Upper Sarmatian (ca. 9 Ma) that are attributed to the Ponto-Caspian gammarids (details, including priors, provided in Copilaş-Ciocianu et al., 2019). The dataset for analysis was supplemented with intermediate lineages between Ponto-Caspian gammarids and Ohrid complex belonging to other species of gammarids that have both COI and 28S markers (data from Copilaş-Ciocianu et al. 2023, Hupało et al. 2020, Mamos et al. 2016, Wysocka et al. 2014). Data were divided into four partitions: all three COI codons and 28S rDNA, following the partitioning scheme selected by PartitionFinder 2.1.1 (Lanfear et al., 2012), evolutionary models were set through bModeltest. The relaxed clock and Birth-Death tree were selected as priors through the AICM method of moments estimator (Baele et al. 2013). Four MCMCM runs, each with 60,000,000 generations and a sample frequency every 3,000 generations were applied in BEAST. All runs and parameters were examined and proved to have ESS>200. The runs were combined, and final trees annotated as above. In order to obtain COI substitution rate for the COI marker the above analysis was also run with codon positions as on partition. Additionally, we estimated substitution rates using as calibration hypothetical age for the radiation of the endemic species-flock in Lake Ohrid. According to the latest report on the deep drilling project from Lake Ohrid (Wilke et al., 2020), deep-water conditions were established ca. 1.2 Ma with the beginning of the Lake 1.36 Ma. Therefore, we use these dates as the constraints for the radiation of the lacustrine flock. For the analysis, LogNormal distribution constraints were set for the most common ancestor of flock species with M: 0.18, S: 0.079 and a median of 1.2 Ma and 95% HPP 1.36-1.05, capturing the distribution with the establishment of the lake with shallow water conditions. The priors (tree and substitution model), MCMC settings and tree processing were done as above.

To explore if COI in species-flock is subjected to positive selection that may deviate its substitution rate from the general one of the *Gammarus balcanicus* species-group, we calculated rates of nonsynonymous (dN) to synonymous (dS) mutations. The *dNdS::ape* function (Paradis & Schliep, 2019) was used, based on Li’s, (1993) proposition of unbiased estimation of rate of dN to dS. As the two groups for comparison we used (*i*) all haplotypes of the *Gammarus* species-flock of Lake Ohrid and (*ii*) all haplotypes of the phylogenetically closest sister lineages (*GcfBAL1-16* and *Gammarus dulensis*; GenBank accession nos: JX899174-77, KJ462705-15, KJ462727-35, KJ462754), published by Wysocka et al. (2014). To test for significant differences between the rates obtained for (*i*) and (*ii*), we run a t-test. To ensure that data passes the test of differences in variances, we run a Fisher’s F-test. For the results suggesting that groups are different (p<0.05, heteroscedasticity), we run a Welch t-test. All statistical analyses were performed in R.

### 2.6 Demographic analysis

Demographic analyses were run for each MOTU with 10 or more representatives. The historical demographic patterns were explored using two approaches. First, to test for a recent demographic expansion, Tajima’s D, Fu’s Fs and Ramos-Onsins and Rozas R2 were calculated using DnaSP 6. Their statistical significance was evaluated using coalescent simulations with 1,000 replications. Additionally, to visualise possible demographic events (i.e. through “star like” topologies), median-joining (MJ) networks (Bandelt et al., 1999) were constructed using PopART 1.7 (Leigh & Bryant, 2015) with homoplasy level parameter set at the default value (ε = 0).

Second, extended Bayesian skyline plots (eBSP) in BEAST 2.6.2 were used to visualise demographic changes through time. The substitution model was set through bModelTest. The molecular clock calibration utilises COI substitution rate obtained from calibration based on geological time of the lake formation (see above). The population model was set to 0.5 and prior on population mean was set to normal distribution with M=1 and SD=0.1. Two MCMC chains were run to ensure convergence. Each was 40 M iterations long, sampled every 20,000 iterations for eBSP logger. One run for each data set was used to plot the eBSP in an R script, applying a 10% burn-in phase.

## 3. Results

### 3.1 Molecular species determination and diversity

Altogether, we obtained 600 COI sequences that, after trimming, collapsed into 85 haplotypes representing all the morphospecies known from Lake Ohrid and 40 sequences of 28S rDNA, at least one for each COI MOTU. Twelve MOTUs of the local *Gammarus* species-flock, inhabiting lake and springs on its shoreline, were identified, by BINs, barcoding-gap-based method ASAP and bPTP (Fig. 2, Fig. S1.1); the mPTP showed 11 MOTU’s, collapsing two newly discovered lineages. MOTUs were named in relation to their morphological affinity, using the following abbreviations throughout: SKE (ACH8452) - *G. sketi*, SOL (ACI1902) - *G. solidus*, CPAR (ACH5405) - *G. cryptoparechiniformis*, PAR_N (AEO6047) - *G. cf. parechiniformis*, CSAL (ACH5669) - *G. cryptosalemaai*, SAL (ACI16561) - *G. salemaai*, SAL_N (AEW2637) - G. cf. salemaai, LYCH (ACH5404) - *G. lychnidensis*, PAR (ACI1559) - *G. parechiniformis*, LYCH-PAR-OCH (ACH8755) - MOTU containing individuals affiliated morphologically to *G. lychnidensis*, *G. parechiniformis* and *G. ochridensis*, STA-LYCH (ACH8754) - MOTU containing morphospecies *G. stankokaramani* and G. *lychnidensis*, MAC - *G. macedonicus*, SYW - *G. sywulai.* PAR_N and SAL_N are newly discovered MOTU showing morphological affinities to *G. parechiniformis* and G. salemaai, respectively. PAR_N contains only three individuals, sharing the same haplotype. The LYCH,bvMAC and SAL_N were represented by single individuals, while the MOTUs STA-LYCH and LYCH-PAR-OCH were the most abundant and the richest in haplotypes (Fig. 2). Both new COI MOTUs (PAR_N and SAL_N) have also private 28S haplotypes, while MOTU MAC share 28S haplotype with STA-LYCH and LYCH with LYCH-PAR-OCH (Fig. 2b). The molecular distance within the *Gammarus* species-flock in Lake Ohrid is highest between SYW and SOL (0.080 p-distance, 0.085 K2p) and the lowest between STA-LYCH and LYCH-PAR-OCH (0.021 p distance, 0.022 K2p). The molecular distance between the *Gammarus* species-flock in Lake Ohrid and its sister lineage SKE ranges between 0.087 and 0.104 for p-distance, and between 0.093 and 0.113 forK2p distance (Table S2.2).

**Fig. 2.**
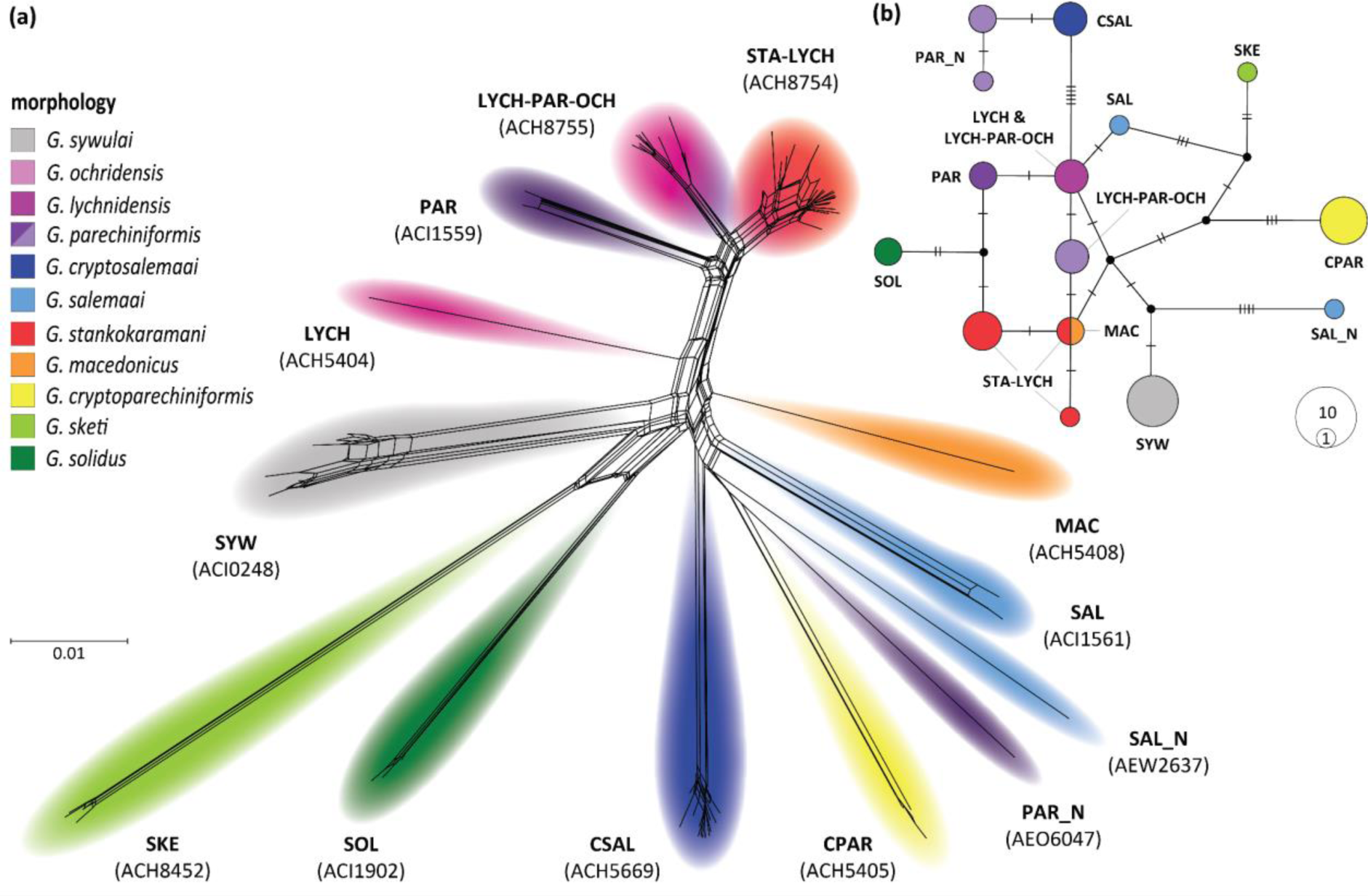
Haplotype relationships of COI and 28S markers. Colours represent morphospecies. MOTUs names (details in Results) and their respective morphospecies. (a) Phylogenetic network of the *Gammarus* species-flock and its sister lineage (*G. sketi*), reconstructed using p-distance between COI haplotypes; (b) TCS haplotype network of 28S rRNA marker. Black dots indicate median vectors (ancestral, unsampled or extinct haplotypes). Lines at the branches denote the number of mutational steps between haplotypes. Circle sizes are proportional to the haplotype frequency.

### 3.2 Spatial and depth-related diversity patterns

The flocks’ sister species (*G. sketi*), representing a single MOTU (SKE), was found exclusively in springs adjacent to Lake Ohrid at its southern and eastern sides: the Biljana springs in Ohrid town, the entire St. Naum springs complex, and a small spring area in Albania near the Albanian-Macedonian border (Fig. 1, Fig. 3). Another spring representative, *G. cryptosalemaai* (CSAL), was found in the latter locality and on the shores of the lake. The spring MOTU PAR representing the morphospecies *G. parechiniformis* has a wider distribution range than SKE, including also the Drilon spring in Albania (Fig. 3); a single specimen was found also on the shoreline of Ohrid, next to St. Naum spring. The MOTU LYCH-PAR-OCH has the largest distribution range with respect to habitat zones in the lake, and can also be found in springs, where only morphotype of *G. parechiniformis* from this MOTU can be found. This MOTU is also present in all habitat zones, reaching the shallowest part of profundal (Fig. 4). PAR_N was found from the littoral to shallow profundal. So far, CPAR has the most narrow range and was found only in the littoral zone, same as LYCH and MAC of which only singular individuals were found in sublittoral. Multiple specimens of SYW were found only in a narrow range of sublittoral, mostly in the *Chara*-belt. The ranges of SOL and STA-LYCH extend from the upper sublittoral to the deepest profundal at over 290 m. In result, sublittoral is the zone accommodating the highest richness of endemic *Gammarus* in Lake Ohrid (8 MOTUs). The littoral and the profundal zones are each inhabited by 4 MOTUs (Fig. 3, 4). Additionally, the sublittoral zone holds the highest haplotypic (H_d_) and nucleotide diversities (π), followed by springs and the profundal zone (Table S2.3).

**Fig. 3.**
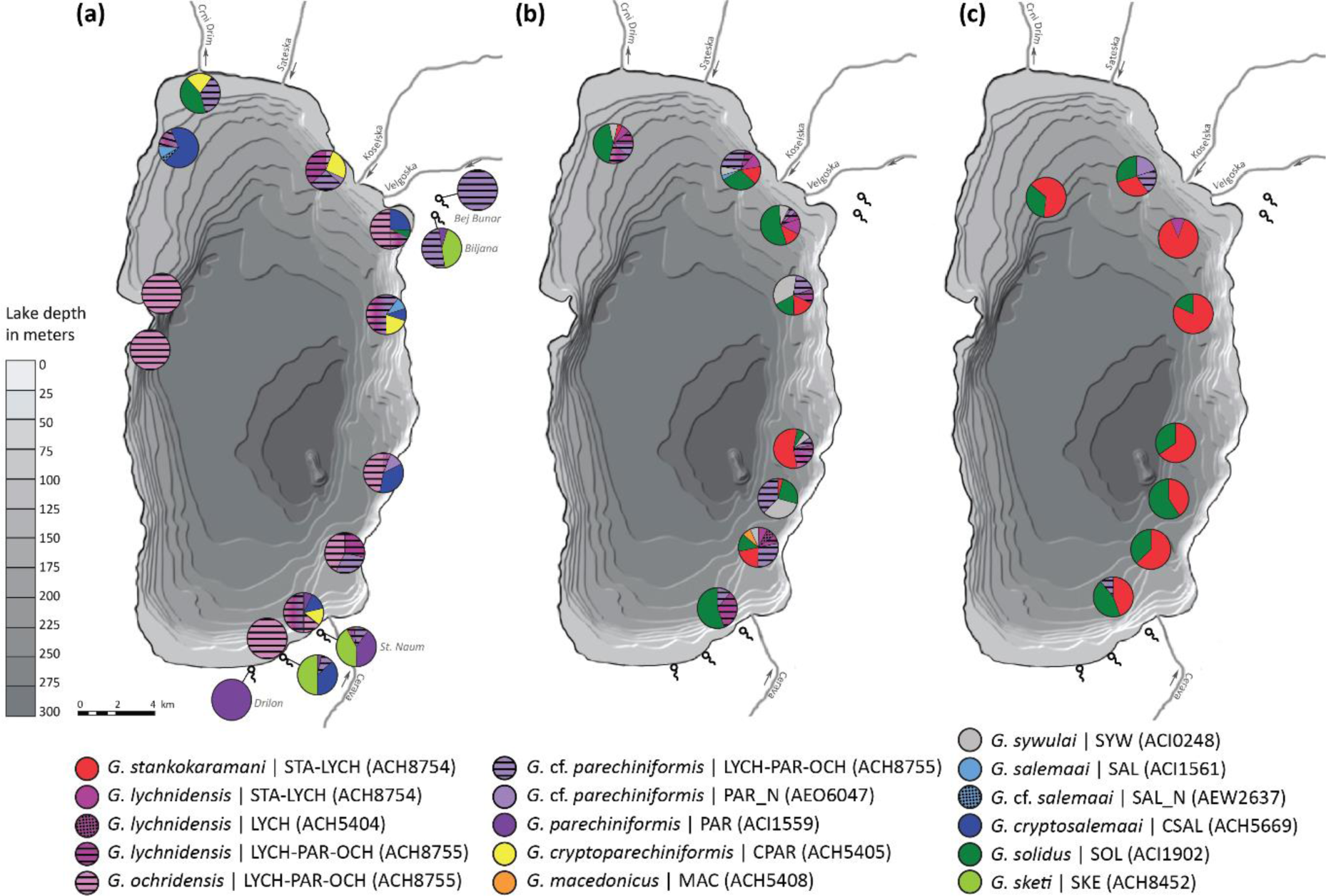
Horizontal distribution of Lake Ohrid *Gammarus* species-flock and its sister species *G. sketi*. Pie charts show relative frequency of morphospecies/MOTUs in sampling sites, colours of pies represent morphological species, white stands for lack of morphological determination, springs marked with a symbol. Maps present different habitat zones: (a) springs and littoral (0-20 m), (b) sublittoral (20-50 m), (c) profundal (50-290 m). Background map modified from Reed et al. (2010).

**Fig. 4.**
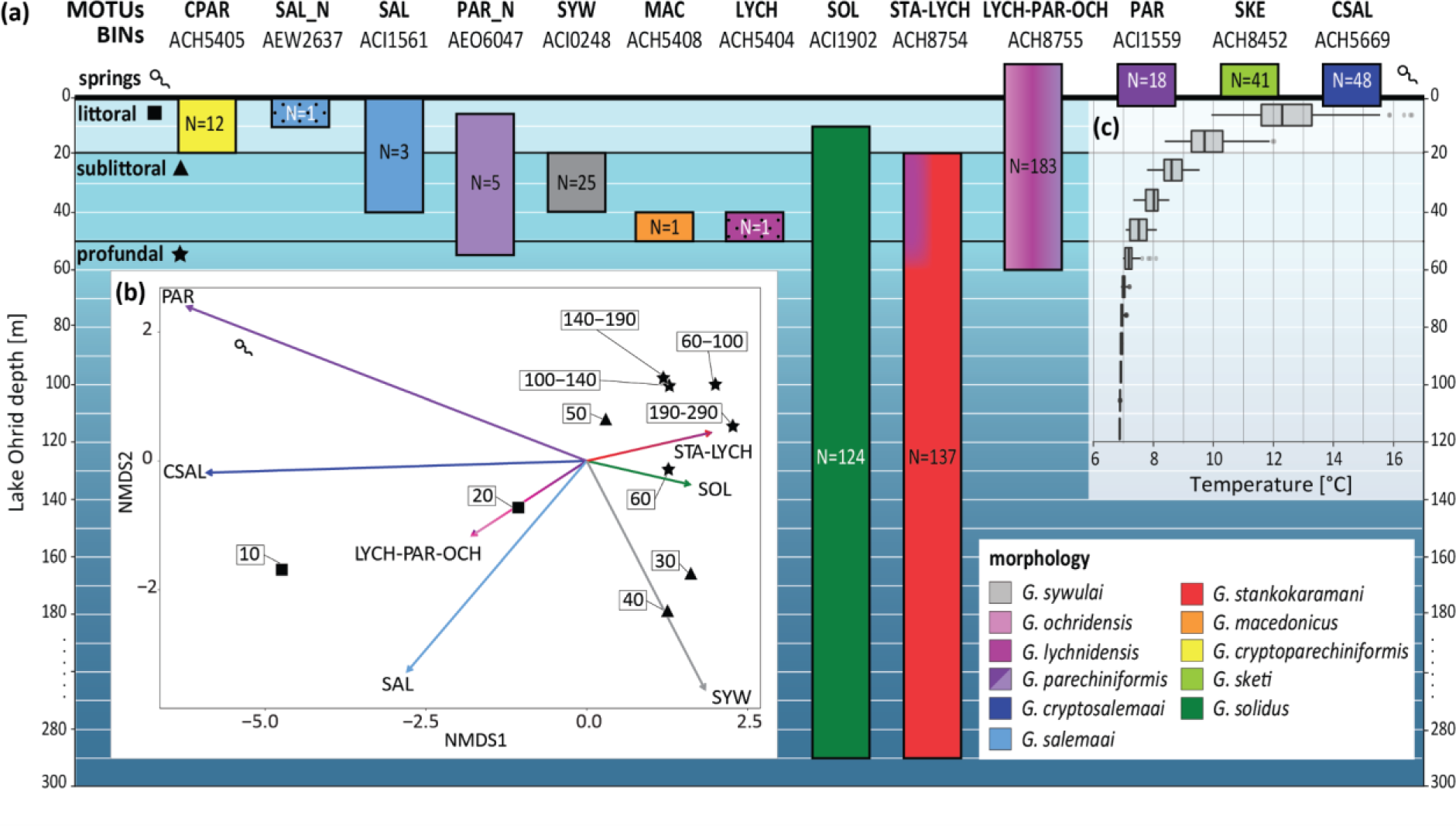
Vertical distribution and diversity of Lake Ohrid Gammarus species-flock and its sister species *G. sketi*. (a) Vertical ranges, colours represent morphological identifications, white stands for lack of morphological determination, N - number of individuals, dashed rectangles enclose spring populations according to MOTU occurrence in each spring system (compare Fig. 5). (b) NMDS analysis of communities grouped for every 10 metres of depth (provided in boxes). (c) Box plots of temperatures reported every 10 metres in the lake down to 120 m.

Considering environmental parameters, the results of the Spearman correlation analysis show that there is a strong positive correlation between the mean and SD of temperature and relative salinity and the number of MOTUs, while all of them are negatively correlated with depth. Also, the Pearson correlation analyses revealed significant correlations in most comparisons, but not for depth *versus* mean temperature, number of MOTUs *versus* mean temperature, nor for SD of temperature and salinity (Table S2.4).

The analysis of variance (permutational MANOVA) performed, both, using the three habitat zones in the lake and, additionally, using the three zones and coastal springs as a fourth one, showed that they significantly structure the composition of *Gammarus* communities in all the analysed cases (Table S2.5). The NMDS analysis suggests that MOTUs: LYCH-PAR-OCH, STA-LYCH, PAR, SOL, CSAL, SAL and SYW have the greatest impact on the distribution pattern of communities (Fig. 4b, Table S2.5). The ordination shows clear distinction of spring communities, separation of litoral and depths between 20 and 40 m of sublittoral. The sublittoral at the depth of 40-50 m, shows close affinity to profundal housing a very uniform community.

### 3.3 Population structure

Analysis of molecular variance showed that, in the case of MOTU LYCH-PAR-OCH, the grouping of sampling sites according to their lacustrine *versus* spring origin explained 53% of the observed variance (Table S2.6). The F_st_ between the two groups has a value of 0.54 (p<0.05), indicating significant differences in their genetic composition. No spatial genetic structure related to habitat zones was detected for the two other abundant and widely distributed MOTUs, STA-LYCH, SOLs (Table S2.6). On the other hand, MIGRATE analysis suggested that the models assuming division of the population to habitat zones is better than the one assuming panmictic populations for LYCH-PAR-OCH (by 45 BF), STA-LYCH (by 30 BF) and SOL (by 6 BF) (Table S2.7). No evident unidirectional gene flow could be detected in the case of STA-LYCH and SOL, and only partially unidirectional gene flow from the lake to springs was indicated in the case of LYCH-PAR-OCH.

### 3.4 Phylogeny and age of the *Gammarus* species-flock in Lake Ohrid

According to our phylogenetic reconstructions based on the COI marker, the *Gammarus* species-flock in Lake Ohrid is monophyletic group, sister to the *G. sketi (SKE)*, and including *G. cryptosalemaai* (CSAL), *G. parechiniformis* (PAR), as well as the MOTU LYCH-PAR-OCH, all three found both in springs and in the lake. The 28S rDNA phylogeny clusters all the flock species together with SKE into one monophyletic clade. Trees based on both markers do not show bigger conflicts in lineages relationships, excluding MAC being nested in STA-LYCH clade and LYCH in LYCH-PAR-OCH in 28S rDNA phylogeny. However, both reconstructions are largely unresolved (Fig. 5a).The phylogenetic reconstructions show also that the newly discovered MOTU SAL_N diverged early during the flock diversification and its molecular affinity to other MOTUs is uncertain. The other new MOTU, PAR_N, clusters with CSAL in case of 28S rDNA reconstruction but shows no affinities within the flock to other MOTUs in case of COI phylogeny.

Based on the molecular clock reconstruction of phylogeny (Fig. 5b, Fig. S1.2), the *Gammarus* species-flock in Lake Ohrid started to diversify ca. 2 Ma (95%HPD: 0.7-4.5 Ma) and the youngest MOTU diverged at ca. 0.5 Ma (95%HPD: 0.1-1.1 Ma). The substitution rate for COI obtained using the geological calibration point (0.0487 My^-1^) is much higher than the one derived from the fossils of Ponto-Caspian amphipods (0.0197 My^-1^, Table S2.8). The rates of nonsynonymous to synonymous mutations (dN/dS) are generally higher in the *Gammarus* species-flock in Lake Ohrid than in outgroups with the means of 0.029 and 0.014, respectively (Table S2.9, Fig S3). The Welsh two sample t-test support significant difference between flock and outgroup dNdS rates (p <0.05, Table S2.9).

**Fig. 5.**
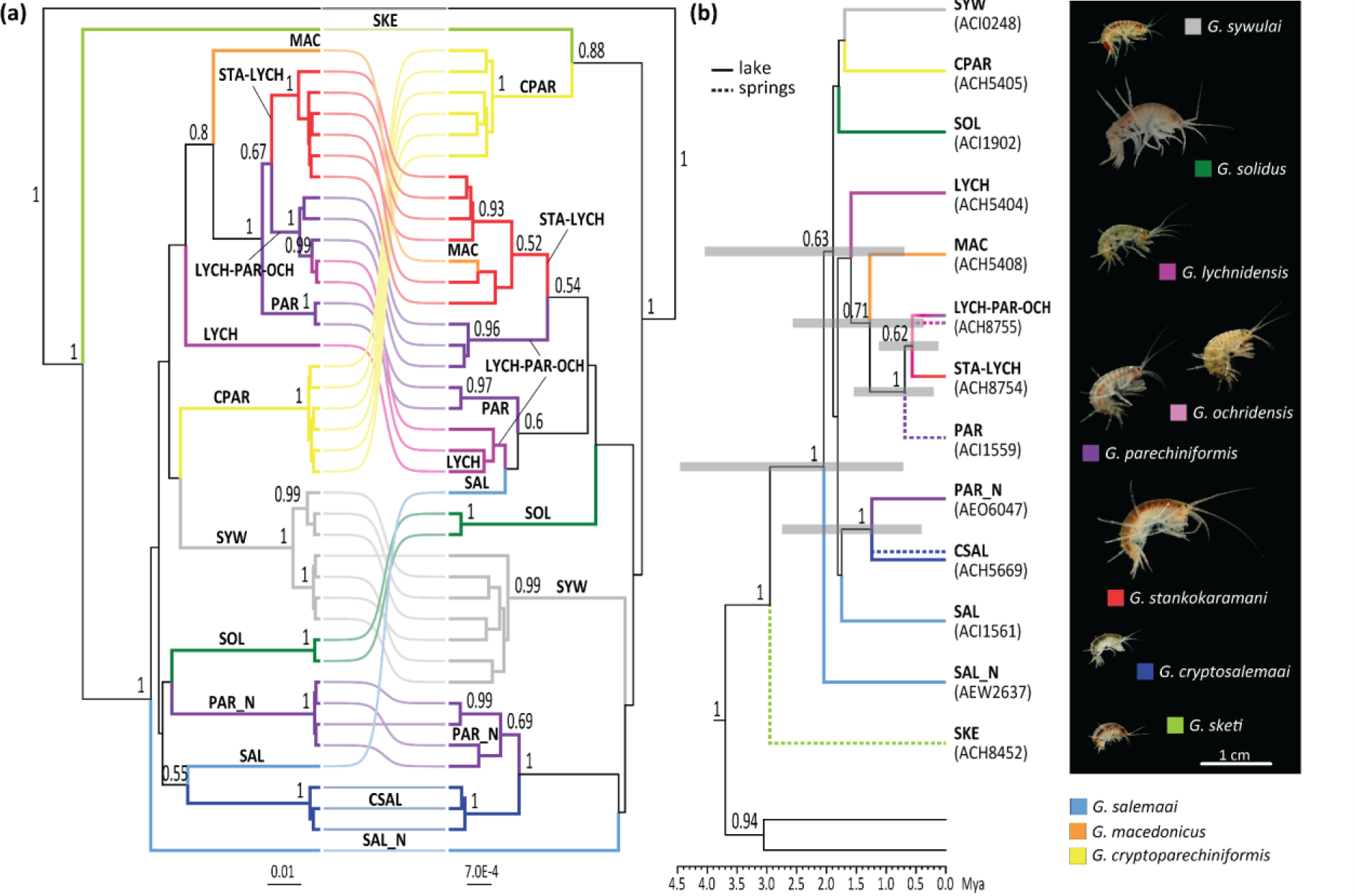
Phylogeny and age of the Gammarus species-flock in Lake Ohrid. Number on nodes stands for posterior probabilities above 0.5. (a) Co-phylo plot depicting Bayesian phylogeny reconstruction based on COI (left) and 28S rRNA (right) markers. *Gammarus* cf. *balcanicus* used as an outgroup. (b) Time calibrated Bayesian reconstruction of phylogeny based on COI and 28S with one specimen per MOTU. Colours represent morphological species. Grey bars represent 95% highest probabilities density range. On the right, photos of Gammarus morphospecies from Lake Ohrid and adjacent springs.

### 3.5 Demographic history of the flock

The results of eBSP suggest that, except PAR, all MOTUs went through some changes of effective population size (Table S2.10). In the case of CSAL, LYCH-STA, SOL and SKE the 95% HPD excludes 0, rejecting a constant population size for these MOTUs during their diversification. In the case of SYW the results suggest that this species had the most stable population size of all the MOTUs. The spring MOTUs, SKE and PAR show signs of rather recent demographic increase (eBSP) after putative bottlenecks (Tajima’s D, Fu’s F). The spring and lacustrine MOTU CSAL show clear signs of population increment 12-15 kya after a putative bottleneck. The lacustrine lineages, LYCH-PAR-OCH, STA-LYCH, SOL and SYW, show a demographic expansion with statistical support for a putative bottleneck predating their expansion. MOTU CPAR does not show signs of demographic changes except for Tajima’s D. The suggested time of these demographic changes varies depending on the two calibration schemes employed. The one based on the geological calibration point (Fig. 6) suggests that the population growths are very recent (last ky); in the case of LYCH-PAR-OCH and CSAL the demographic growth is suggested to happen at the beginning of Holocene.

**Fig. 6.**
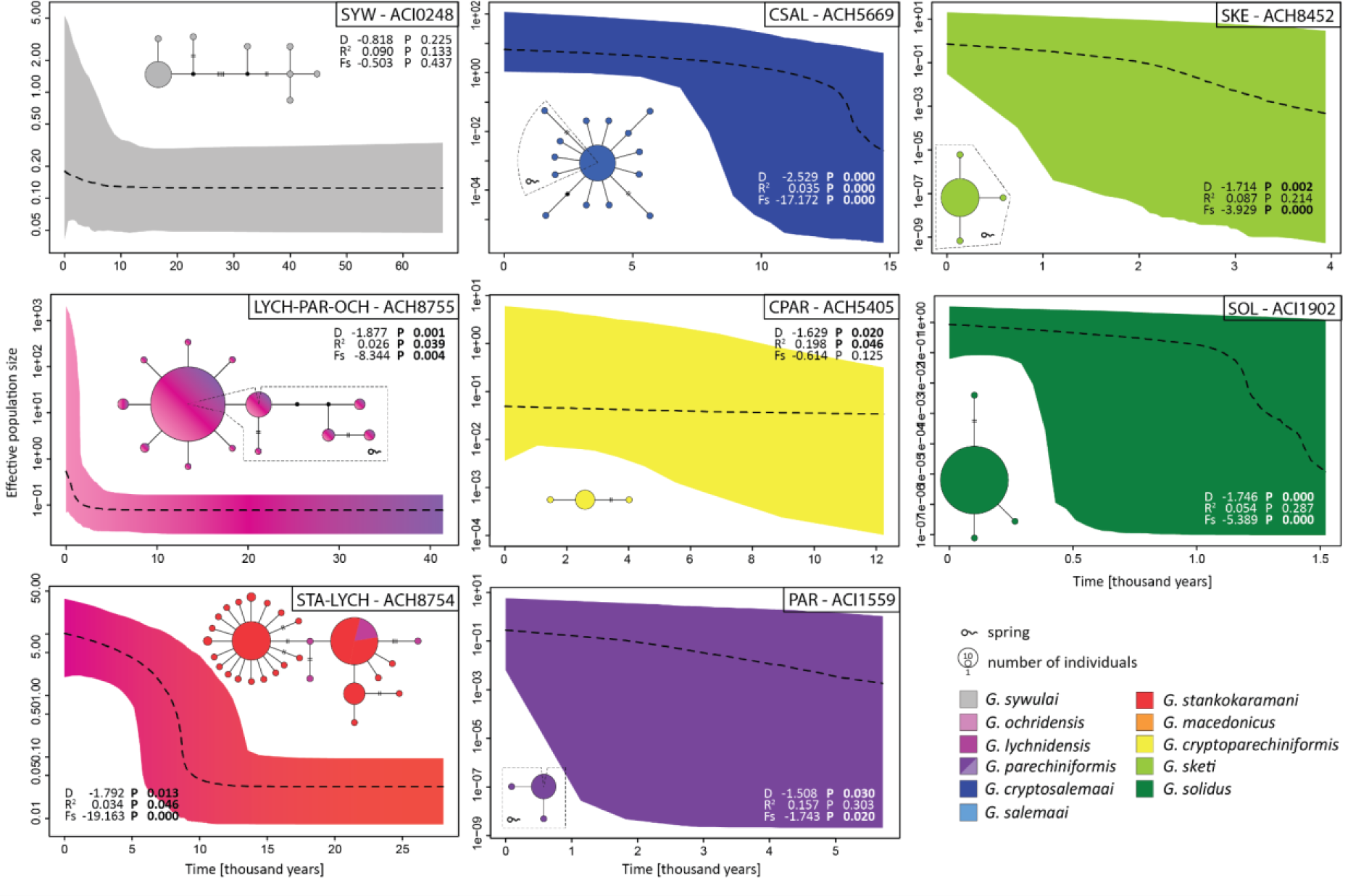
Demographic history based on the COI marker of the MOTUs belonging to the *Gammarus* species-flock from Lake Ohrid. Dating is based on the calibration point set for establishment of deep water conditions in the lake (see Materials and Methods). Analysis was performed for MOTUs containing 12 or more individuals. Plots of eBSP and haplotype networks with MOTU code above, attributed with results of neutrality tests (Tajima’s D, Fu’s Fs, Ramos-Onsins and Rozas R2) and their respective p values.

## 4. Discussion

### 4.1 Diversity and conservation

The *Gammarus* species-flock inhabiting ancient Lake Ohrid is exceptional on a global scale and unique for the freshwater genus *Gammarus*. The flock belongs to the *G. balcanicus* species group found predominantly in the submontane and mountain streams and springs of central, eastern and southern Europe (Wysocka et al., 2014). Until now, eight MOTUs from Lake Ohrid and two from the neighbouring springs, the latter sympatric with *G. sketi*, were known (Grabowski et al., 2017). In the presented study, through DNA barcoding, supported by the nuclear 28S RNA marker, two new intralacustrine MOTUs (SAL_N, PAR_N) were recovered and a range of two, previously considered as exclusively spring MOTUs (CSAL, PAR), were extended to shorelines of the lake. Taking into account the molecular distance of the new MOTUs to the rest of the flock and phylogenetic position, they likely represent two species new to science. Surprisingly, despite wider sampling, the MOTUs LYCH and MAC were not recovered in our study. So far, the MOTU MAC is the only representation of the species *G. macedonicus*. These two MOTUs are represented by single individuals collected only during 2007-2011 samplings (Wysocka et al., 2013). Moreover, the MOTU SAL (*G. salemaai*), represented by three specimens, was also not detected since these past sampling campaigns. Recording of two new MOTUs points out to the importance and suitability of DNA barcoding for the illumination of biodiversity, even in theoretically well-studied water reservoirs. On the other hand, the lack of recovery of any specimens belonging to the above-mentioned three MOTUs, despite extensive sampling, already suggests a possible loss of diversity, happening over the last decade.

Such putative loss of diversity is disturbing and may be the consequence of multiple factors that influence degradation of Lake Ohrid. These include extensive urbanisation and tourism development, both increasing pollution of the lake and destruction of spring habitats (Zdraveski et al., 2021), and/or introduction of invasive species (Trajanovski et al., 2019). Climate change is another factor that might contribute to the loss of endemic diversity in Lake Ohrid (Matzinger, 2006, 2007). Lake Ohrid is expected to experience a decreased vertical mixing and less frequent deep convective mixing as global temperatures increase. Over time, this is predicted to interplay with eutrophication and nutrient inputs, contributing to decline in dissolved oxygen levels, particularly in deep water habitats, impacting endemic fauna (Matzinger et al., 2007). Taking into account that 90% of the amphipod fauna in Lake Ohrid is endemic and that the majority of other organisms is also endemic and often relict (summary in Albrecht & Wilke, 2008), conservation efforts to save this unique ecosystem are sorely needed.

### 4.2 Distribution

The sister species of the Lake Ohrid *Gammarus* species-flock - *G. sketi*, represented by a single MOTU (SKE) was originally described from the Biljana springs in the town of Ohrid (Karaman, 1989) and found nowhere else. Later studies (Wysocka et al., 2013) recovered it only in the “southern” springs in Albania and in St. Naum. Our current work confirms its presence in Biljana and in the other springs. Its distribution, together with *G. parechiniformis* (MOTU PAR) suggests interconnectivity of the springs and underground dispersal capabilities of these gammarids. Such a subterranean hydrological connection is already known between Lakes Prespa and Lake Ohrid, including its springs (Amataj et al., 2007). Already, in the original description of the species, Karaman (1989) reported *G. sketi* to have reduced eyes as an adaptation to subterranean life.

Our study does not show any clear patterns in horizontal distribution of morphospecies and MOTUs. However, particularly high diversity was found in areas in the north-eastern and south-eastern parts of the lake, caused by the presence of MOTUs with low numbers of recovered specimens. The south-eastern *Gammarus* “diversity hotspot” corresponds in locality to the Gastropoda biodiversity hotspots proposed by Hauffe et al. (2011). On the other hand, there is a clear vertical differentiation of the flock species, particularly evident in four MOTUs, i.e. CPAR and CSAL found only in littoral *versus* SYW, MAC and LYCH found only in sublittoral. The same bathymetric segregation is observed in the case of Lake Ohrid endemic gastropods. For example, strong vertical segregation is reported for *Ginaia munda munda* (Sturany, 1894) and *G. munda sublitoralis* Radoman, 1978 occupying the littoral (*Chara*-belt) and the sublittoral (shell zone), respectively (Albrecht & Wilke, 2008). A similar division is found for *Ancylus* spp., and *Acroloxus* spp. (Albrecht et al., 2006; Hubendick, 1960). Also, in case of other crustaceans from Lake Ohrid e.g., *Proasellus remyi*, such a vertical segregation is observed between three morphological forms (f*. remyi,* f. *acutangulus,* f. *nudus*). Where *P. remyi* f*. remyi* dominates in littoral, *P. remyi* f*. acutangulus* dominates in sublittoral, where both forms partially overlap (Wysocka et al., 2008). In the case of the *Gammarus* species-flock, the MOTUs SYW, PAR_N and LYCH-PAR-OCH overlap in the littoral and the sublittoral, with the latter two reaching profundal. However, this partially profundal distribution is probably an artefact as the habitat zones do not always have sharp boundaries at the same depth throughout the lake, and the MOTUs are associated with the *Chara*, rocks and shell zones. In contrary to the isopod, *P. remyi* f*. nudus* - restricted to profundal, the two deep water species, *G. stankokramani* (STA-LYCH) and *G. solidus* (SOL), extend their ranges up to the border between sublittoral and littoral. Such a pattern of gammarid distribution, with some species predominating in littoral and sublittoral and some that are eurybathic and found in depths of hundreds of metres, but also in sublittoral, is known from Lake Baikal, the most species-rich radiation of gammarids in any ancient lakes (Takhteev, 2000) and partially also from the Ponto-Caspian basin (Copilaș-Cioicianu & Sidorov, 2022). Interestingly, according to our study, depth, temperature and relative salinity correlate to vertical diversity, suggesting that these physical factors may play a role in structuring gammarids diversity. Even standard deviation of temperature and salinity is correlated to each of the physical factors, which is not surprising as these parameters are getting more stable and diversity decreases as the depth increases.

According to the MIGRATE analysis, the model with the depth-associated separation of populations outperforms the panmictic one, supporting the bathymetric segregation according to habitat zones at the MOTU level in case of LYCH-PAR-OCH, SOL and STA. However, there is no clear pattern of gene flow between the habitat zones, suggesting symmetric migration. AMOVA did not show any spatial population structure, except for LYCH-PAR-OCH lake *versus* spring populations. The MOTU LYCH-PAR-OCH, distributed both in lake and neighbouring springs, is a peculiar case, as it contains three morphospecies: *G. lychnidensis*, *G. ochridensis and G. parechniformis*. Only the latter morphospecies can be found in springs. According to phylogenetic analysis and the occurrence of private haplotypes in the Bijlana springs, the MOTU probably has a spring origin. After the colonisation of the lake from the springs of Biljana, it probably started to colonise the southern springs, including St. Naum, as can be concluded from the putative direction of gene flow (spring to lake according to MIGRATE) and the fact that only the haplotype widely distributed in the lake can be found in these springs.

### 4.3 Drivers of diversification

The differences in vertical distribution of MOTUs and patterns of MOTU populations segregation to habitat zones strongly suggest that parapatric speciation is the likely mode of diversification in the *Gammarus* species-flock of Lake Ohrid. In parapatric speciation, divergence of neighbouring or partially overlapping populations is suggested to happen as a consequence of low gene flow, for example via local adaptation, with the subsequent build-up of reproductive isolation (Coyne & Orr 2004; Futyuma, 2005). Considering the habitat preferences of MOTUs belonging to the *Gammarus* species-flock in Lake Ohrid, speciation indeed seems to proceed along an ecological gradient, as suggested for ancient lakes by Schön & Martens (2004). Given the specific habitat zonation in Lake Ohrid, parapatric speciation was also suggested as a driver of diversification in other animals in Ohrid (summary in Albrecht & Wilke, 2008).

Moreover, the substantial vertical and horizontal overlap of morphospecies and MOTUs in Lake Ohrid gammarids, the rapid accumulation of lineages at an early stage of their diversification and the decoupling of morphological and molecular diversity, suggests that gammarids underwent adaptive radiation in Lake Ohrid. Adaptive radiation is the rapid origin of new species from a single ancestor as a consequence of adaptation to distinct environments (Schluter, 2000). However, at the moment we do not have enough information about the flock’s ecological preferences and respective morphological traits to verify the adaptive nature of the gammarid radiation in Lake Ohrid.

### 4.4 Age and the demographic history of the flock

The calibration used in our study suggests that the flock originated ca. 2 Ma (between 0.7 and 4.5 Ma), which is in congruence with the first study utilising molecular dating in investigation of the *Gammarus* species-flock in Lake Ohrid (Wysocka et al., 2013) and partially overlaps with the rate-based radiation shown by Stelbrink et al. (2020). The resulting range also overlaps with the age of establishment of deep water conditions in the lake that happened at ca. 1.2 Ma (Wilke et al., 2020) and with the origin of the lake dated at 1.36 Ma by the SCOPSCO drilling project (Wagner et al., 2017). When using the establishment of deep water conditions as calibration point for radiation of the *Gammarus* species-flock, we obtained a substitution rate of COI faster than the one previously established for the *Gammarus balcanicus* group with independent calibration schemes (Copilaș-Ciocianu & Petrusek, 2017; Mamos et al., 2016) and than the general rate for the Amphipoda, based on fossil calibrations (Copilaș-Ciocianu et al., 2019). The deviation between different calibration schemes and rates can be explained in multiple ways: (*i*) The previous calibration schemes are possibly based on wrong assumptions and, thus, providing incorrect rates. It is highly unlikely, taking account that the calculations were done multiple times, independently utilising different calibration constraints for different arthropods (Brower, 1994). (*ii*) The most recent common ancestor of the flock already existed before the establishment of the lake. Given that crown lineages belong to deep water or predominantly sublittoral organisms, it would imply that the adaptation to lake environment happened multiple times independently. This scenario, however, is not very plausible, given that the flock species phylogenetically nested in the *G. balcanicus* group, known for its preference to well oxygenated small rivers or springs (Karaman & Pinkster, 1987; Mamos et al., 2014) and such innovations as colonisation of a new environment happens rarely. (*iii*) There was an acceleration of COI substitution rate. Substitution rates are known to vary among genes, chromosomes, species, and lineages, which can be due to multiple biological processes. Specifically, among genes, adaptive evolution increases the rate of substitution (Li, 1997). Such increased rates of evolution for specific regulatory or coding sequence, which may relate to substitution rate, has been observed in case of adaptive radiation and interpreted as a signature of positive selection (Brawand et al., 2014). An increased evolutionary rate of COI can be suggested for Lake Ohrid *Gammarus* flock versus sister lineages through a significantly higher ratio of nonsynonymous to synonymous mutations in the former. Such evolutionary accelerations for specific genes are known to occur in other gammarid radiations (Naumenko et al., 2017). Therefore, we tentatively suggest that the diversification of the *Gammarus* species-flock radiation in Lake Ohrid occurred in line with the formation of the lake and resulted in positive selection on *COI* gene and acceleration of substitution rate. However, to properly support this hypothesis, a much larger representation of the genome would be required, preferably full genome sequences with its coding and non-coding parts.

Furthermore, according to our molecular dating, the range of divergence within the flock, especially between MOTU’s found predominantly in springs from those exclusively, or mostly lacustrine overlaps with different interglacials during the Pleistocene when the relative global water levels were fluctuating with high amplitude (Pages, 2016). The high sea water level due to deglaciation probably resulted also in increased precipitation and increase of lakes’ water level like during the last glacial maximum (McGee, 2020). This could have led to inundation of springs located along the shore of Lake Ohrid and divergence of spring ancestor through adaptation to lake conditions. The emergence of new lineages could also have been triggered by hybridisation of spring species with their lacustrine sister lineages. Hybridisation happens in case of species-flocks when lineages diversify rapidly and reproductive barriers are often not completely developed, leading to occasional hybridisation between lineages (Berner & Salzburger, 2015). It is likely that hybridisation stands behind the observed diversity and behind the decoupling of morphological and molecular diversity (Wysocka et al., 2013; Grabowski et al., 2017). This is supported by different patterns of mitochondrial and nuclear divergence (Wysocka et al., 2013) and unique karyotypic patterns. The flock species have chromosome numbers in the range of n=12 to n=34, while in general, gammarids have n=26 (Salemaa & Kamltynov, 1994). The association of water level fluctuations caused by glaciation events in Pleistocene with speciation events for the flock was already proposed as a rather plausible, but not verified, hypothesis (Wysocka et al., 2013).

The putative impact of water level fluctuation can be also observed in case of demographic patterns of particular MOTUs. Especially in the case of STA-LYCH and CSAL, where the population expansion occurs at the beginning of the Holocene. This is often the case of European crustaceans and could be related to hydrological changes with putative increment of freshwater level (Csapó et al., 2020; Mamos et al., 2021; Sworobowicz et al., 2020). In other cases, the demographic changes are more recent, starting at some 3-5 kya and could be a case of colonisations of new depth ranges or springs, like in case of LYCH-PAR-OCH, but are rather hard to interpret. Nevertheless, a recent study of phylogenetic community structure of diatoms from Lake Ohrid showed recent increments in their functional richness and mean phylogenetic distance. These changes are probably associated with variations in water level and affected mixing processes, nutrient content and a set of other potential environmental factors, as diatoms are highly sensitive to changes in the environment (Jovanowska et al., 2022). Moreover, assuming putative positive selection on *COI* gene, we cannot exclude that the low number of haplotypes in case of deep water MOTU SOL and the resulting patterns of recent demographic changes is an effect of selective sweep favouring only specific versions of the gene.

## Conclusions

The endemic Lake Ohrid *Gammarus*-flock show weak horizontal but noticeable vertical differentiation reflecting habitat zonation, with particular MOTUs found predominantly in littoral, sublittoral or profundal. The diversity across bathymetric gradients correlates strongly to temperature and salinity, and shows diversity hotspot in springs and sublittoral. The extensive sampling and DNA barcoding allowed us to detect new MOTU belonging to *Gammarus* species-flock of the ancient Lake Ohrid, representing putatively new species and suggesting that biodiversity of the lake is still not fully discovered. At the same time three other MOTUs escaped recovery which can suggest loss of diversity. In relation to anthropogenic pressure and climate changes, impacting the lake, together with potential loss of diversity, it is clear that conservation efforts to save this unique ecosystem are sorely needed. Altogether, we now know 12 MOTUs endemic to the lake and its spring system. The age of the flock is still uncertain, but most likely coincides with establishment of deep water conditions in Lake Ohrid at 1.2 Ma. The speciation events and demographic changes of the *Gammarus* species-flock are most probably related to glacial and postglacial water level changes, also patterns of very recent population expansions are visible that may be results of colonisation of new depth ranges. Finally, our study proves that, in the world of fast-developing phylogenomic and population genomic methods, DNA barcoding remains a low-cost, handy and reliable tool to provide insight into spatial diversity/local scale biogeography and evolutionary patterns at the species or species-group level in non-model taxonomic groups, for which the detailed genome architecture is unavailable or difficult to obtain for technical reasons (as in gammarids). This may be of particular importance for ancient lakes which hold a variety of invertebrate taxa, presumably many with intricate diversification and speciation histories but unperceived until now. Thus, our approach may provide a useful scheme for the other little studied ancient lakes, such as Ioannina, Doiran, Titicaca or even Tanganyika and Victoria, which are understudied for most of the invertebrate groups.

## Acknowledgements

Authors would like to thank: Dr. Elizabeta Veljanoska Sarafiloska and Dr. Orhideja Tasevska, director of the Hydrobiological Institute-Ohrid for their legal help in organising the fieldwork (permits numbers: 08-237/1, 05-167/1), Zoran Brdarovski, skipper of the institute’s research vessel, for his invaluable help in sampling on Lake Ohrid; Nicolas Boileau, and Fabrizia Ronco from Zoological Institute Basel for help, respectively, in the laboratory and with data analysis in R; Kajetan Deja IOPAN for temperature and salinity data collection; Andrea Desiderato for help with permutational multivariate analysis. This work was supported by the: Polish National Science Centre (OPUS16 grant Nb. UMO-2018/31/B/NZ8/03103); Internal funds of the Faculty of Biology and Environmental Protection, University of Lodz. Tomasz Mamos was supported by the Scholarship of the Polish National Agency for Academic Exchange (NAWA) Bekker Programme (Project Nb. PN/BEK/2018/1/00225).

## Data accessibility statement

All newly generated sequences together with metadata (collection data, GPS codes, taxonomy) are publically available in BOLD dataset DS-OHLGADD (doi provided after acceptance). Additionally, sequences were deposited in NCBI GenBank under accession numbers: (provided after acceptance).

## List of Figures

## Supporting Information

