## Appendix 1 for "Distribution, diversity and diversification from a DNA barcoding perspective: the case of *Gammarus* radiation in Europe’s oldest inland waterbody - the ancient Lake Ohrid"

Fig. S1.1 Barcode gap histogram (molecular distance *vs.* frequency) based on COI generated by ASAP software.

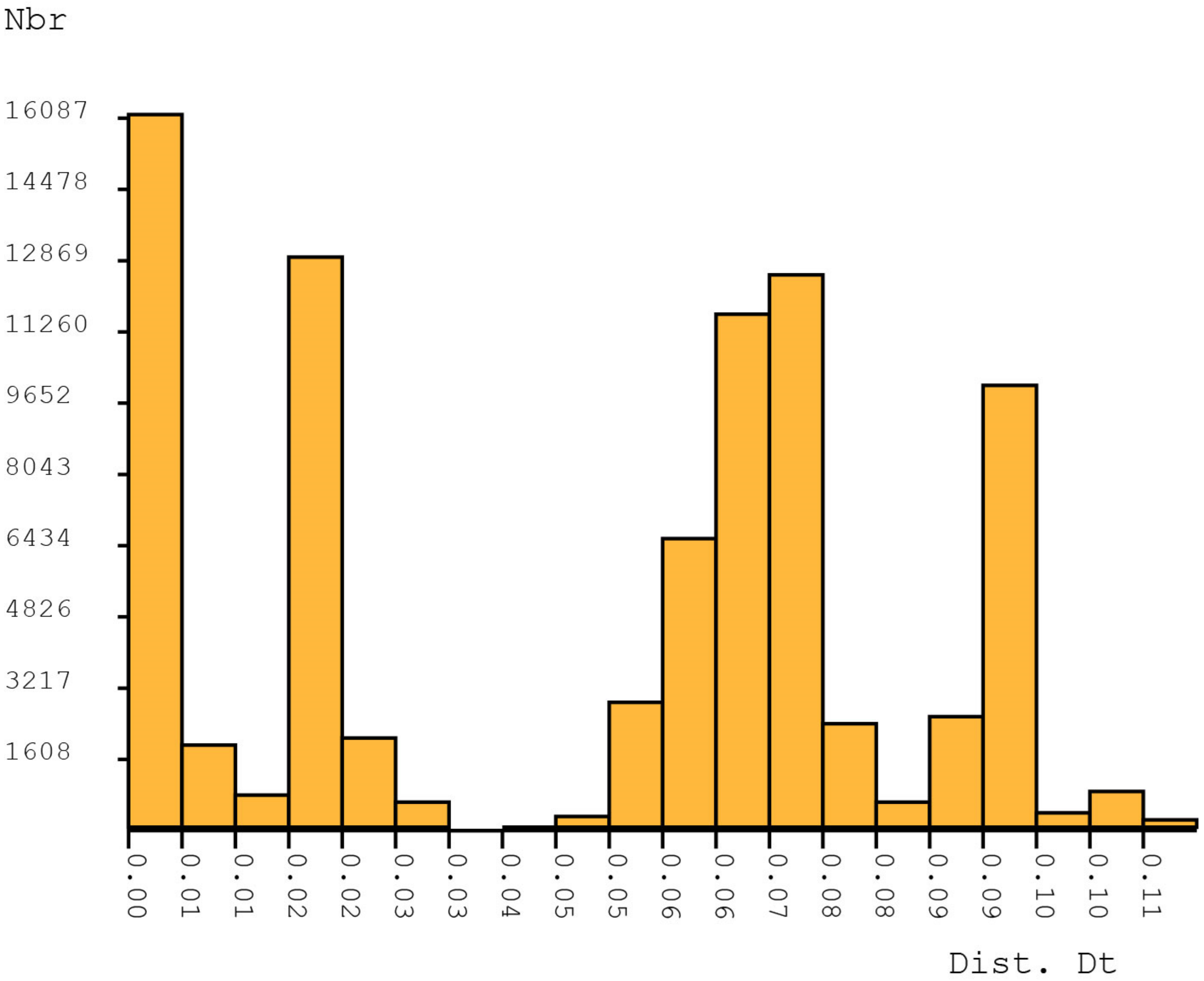

Fig. S1.2 Time calibrated Bayesian reconstruction of phylogeny based on COI and 28S markers. Values on the nodes represent posterior probability. Grey bars represents 95% highest posterior density range. After the pipe symbol are given GanBank codes for COI and 28S sequence, respectively, or BOLD process ID for the specimen. Black circle indicates calibration point.

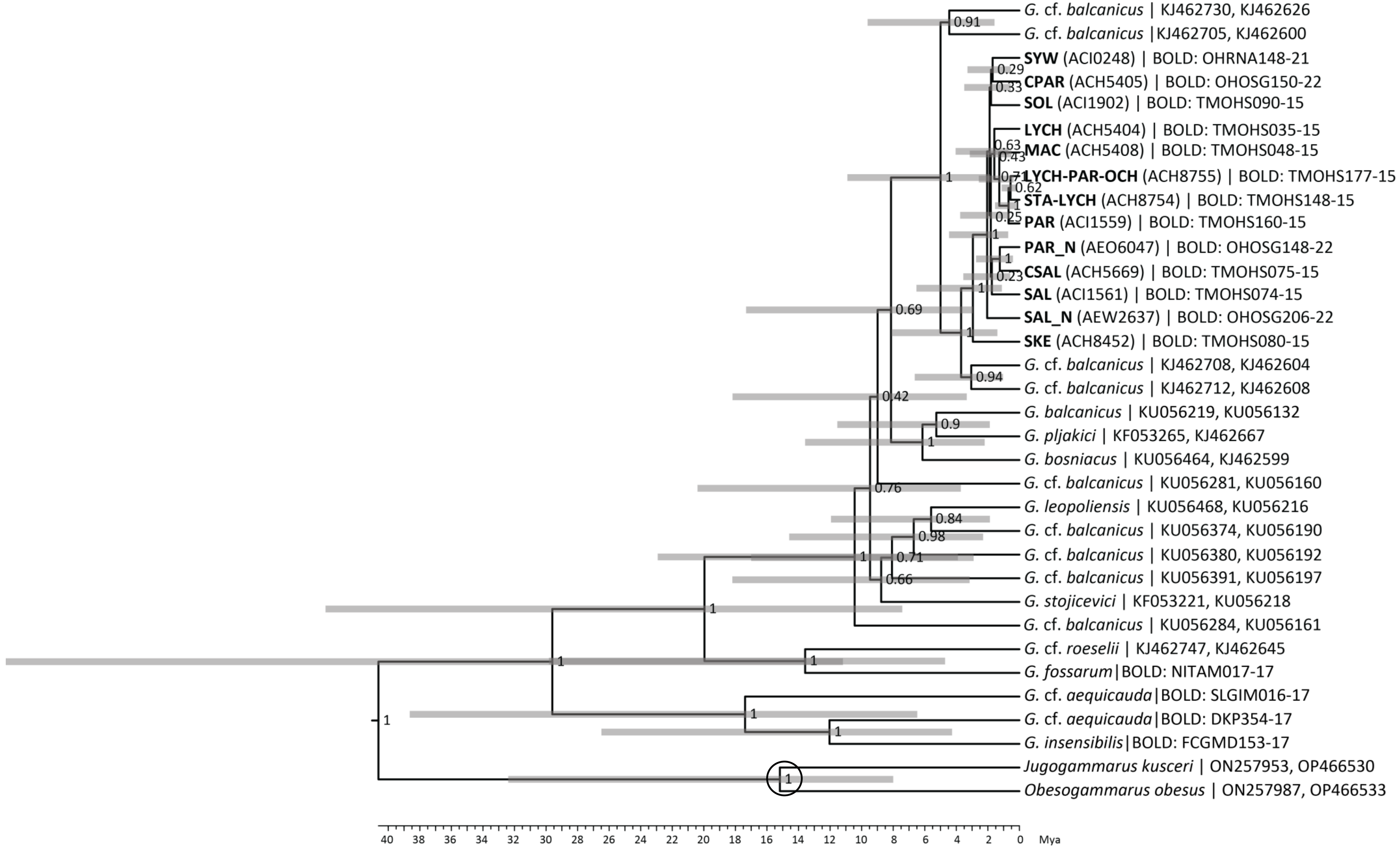

Fig. S1.3 Rates of nonsynonymous (dN) to synonymous (dS) mutations for *Gammarus* species-flock and sister group (details in Materials and Methods).

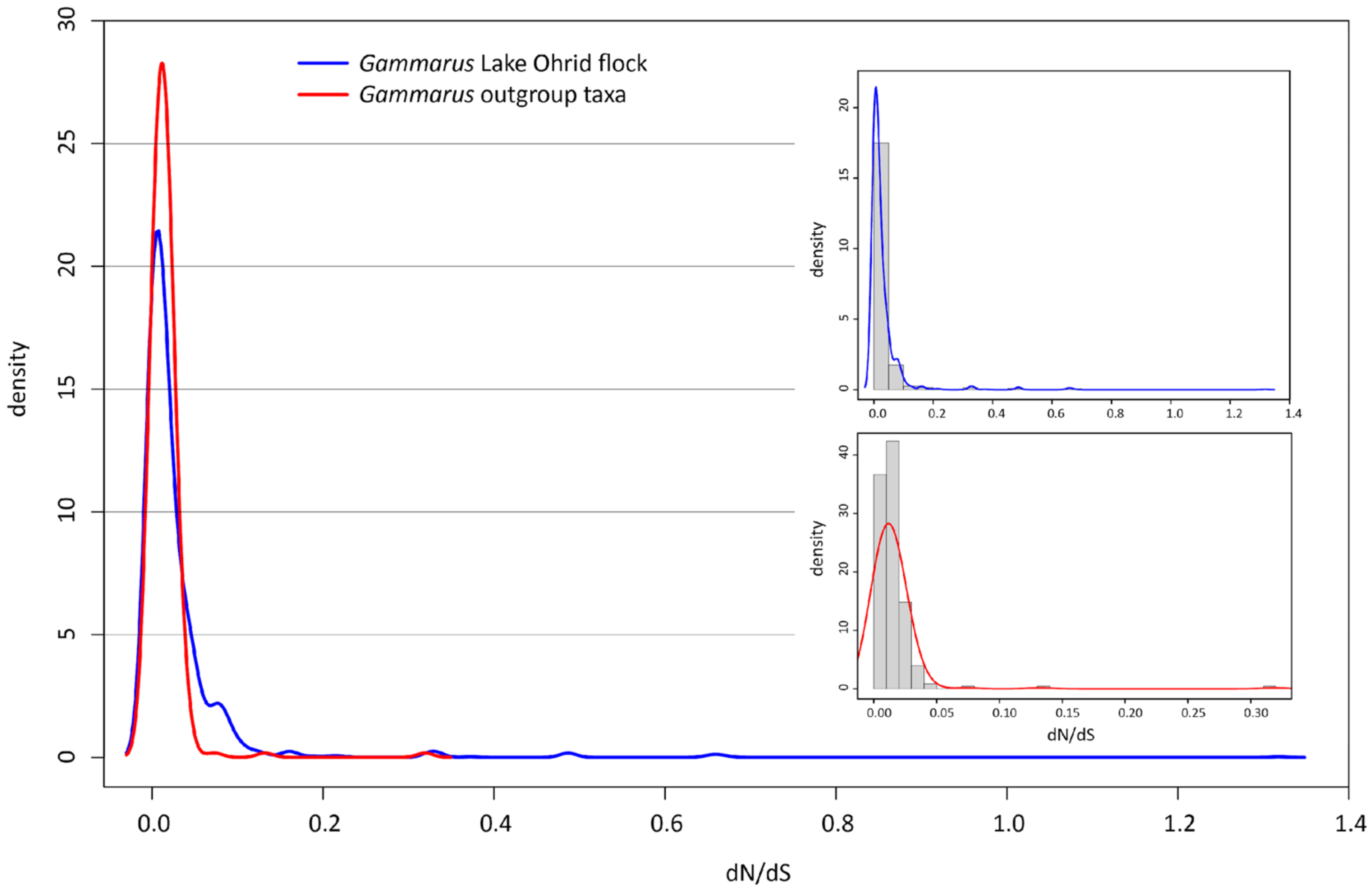
